# Rapid multilocus adaptation of clonal cabbage leaf curl virus populations to *Arabidopsis thaliana*

**DOI:** 10.1101/2021.11.29.468282

**Authors:** J. Steen Hoyer, Olivia W. Wilkins, Aanandi Munshi, Emma Wiese, Divya Dubey, Savannah Renard, Karoline Rosendal Hartø Mortensen, Anna E. Dye, Ignazio Carbone, Siobain Duffy, José Trinidad Ascencio-Ibáñez

**Affiliations:** Rutgers University, Department of Ecology, Evolution, and Natural Resources, New Brunswick NJ, 08901, USA; North Carolina State University, Department of Biological Sciences, Raleigh NC, 27695, USA; North Carolina State University, Department of Molecular and Structural Biochemistry, Raleigh NC, 27695, USA; North Carolina State University, Department of Plant and Microbial Biology, Raleigh NC, 27695, USA; North Carolina State University, Center for Integrated Fungal Research, Department of Entomology and Plant Pathology, Raleigh NC, 27695, USA

**Keywords:** begomovirus mutation, within-host population variability, rapid adaptation

## Abstract

Cabbage leaf curl virus (CabLCV) has a bipartite single-stranded DNA genome and infects the model plant *Arabidopsis thaliana*. CabLCV serves as a model for the genus *Begomovirus*, members of which cause tremendous crop losses worldwide. We have used CabLCV as a model for within-plant virus evolution by inoculating individual plants with infectious clones of either a wild-type or mutagenized version of the CabLCV genome. Consistent with previous reports, detrimental substitutions in the Replication-associated gene (Rep) were readily compensated for by direct reversion and/or alternative mutations. A surprising number of common mutations were detected elsewhere in both viral segments (DNA-A and DNA-B) indicating convergent evolution and suggesting that CabLCV may not be as well adapted to *A. thaliana* as commonly presumed. Consistent with this idea, a spontaneous coat protein variant consistently rose to high allele frequency in susceptible accession Col-0, at a higher rate than in hypersusceptible accession Sei-0. Numerous high-frequency mutations were also detected in a candidate Rep binding site in DNA-B. Our results reinforce the fact that spontaneous mutation of this type of virus occurs rapidly and can change the majority consensus sequence of a within-plant virus population in weeks.

## Introduction

New sequencing technologies have accelerated both virus discovery (Jeske 2018; Cobbin et al. 2021) and studies of within-host virus diversity. Short-read technologies such as Illumina sequencing-by-synthesis provide limited information about viral haplotypes but can yield deep coverage for quantitative analysis of minority (subconsensus) alleles (Lauring 2020). Measuring and functionally characterizing such variation, including spontaneous mutations, is necessary to clarify how virus populations interact with other components of the phytobiome (Roossinck 2019).

Viruses in the family *Geminiviridae* are transmitted by phloem-feeding insects and cause major crop losses (Rojas et al. 2018). Geminivirus populations vary within individual host plants, a fact evident from the first complete-genome sequence dataset (Stanley and Gay 1983). Controlled experiments have demonstrated high geminivirus mutation frequencies (Isnard et al. 1998) and rapid substitution in response to selection (Ge et al. 2007). Geminiviruses have single-stranded DNA genomes yet have substitution rates comparable to RNA viruses (Duffy and Holmes 2008; Duffy et al. 2008). It is thought that rapid evolution enables geminiviruses to adapt and evade host defenses, but the mechanistic details and implications for disease management are largely unclear (Acosta-Leal et al. 2011; García-Arenal and Zerbini 2019).

Cabbage leaf curl virus is a whitefly-transmitted bipartite virus in the genus *Begomovirus* that broadly infects Brassicaceae (Strandberg et al. 1991). We abbreviate the virus name here as ‘CabLCV’, per the Virus Metadata Resource of the International Committee on Taxonomy of Viruses (Calisher et al. 2019), but note that ‘CbLCV’, ‘CLCV’, and ‘CaLCuV’ are also used in the literature (Hill et al. 1998; Hunter et al. 1998; Umaharan et al. 1998; Turnage et al. 2002). The CabLCV DNA-A segment encodes five proteins involved in replication and counterdefense (Figure 1A) and the DNA-B segment encodes two proteins that function in virus movement. CabLCV infects the model plant *Arabidopsis thaliana* (L.) Heynh. (Hill et al. 1998) and dramatically reprograms host gene expression (Ascencio-Ibáñez et al. 2008).

**Figure 1:**
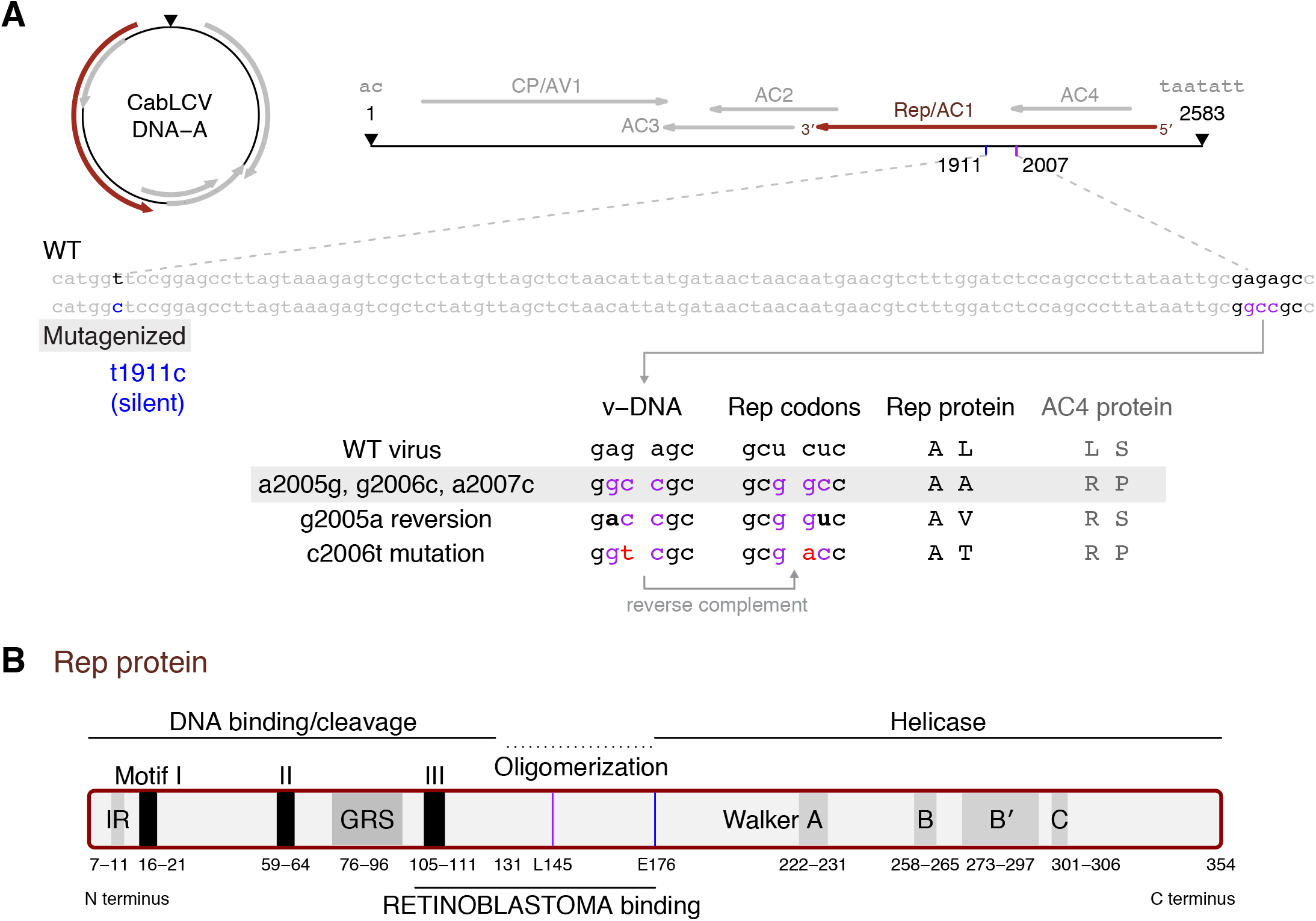
DNA-A mutagenesis and reversion alleles described by Argüello-Astorga et al. (2007). **A.** Circular and linear diagram of the CabLCV DNA-A segment in the conventional orientation, starting and ending at the nick site within the taatatt/ac nonanucleotide (black triangle). The five canonical New World begomovirus genes are indicated, along with the genomic positions of four mutagenized nucleotides in the pNSB1101 plasmid inoculum, in blue (t1911c) or purple (a2005g, g2006c, and a2007c). The sequence display shows the virion-encapsidated strand (v-DNA; v-sense) but note that Rep is encoded on the opposite (complementary) strand. Codons for two amino acid residues in the Rep protein (A144 and L145 for parental wild-type virus) are shown along with amino acid replacements resulting from different mutations; these codons are in complementary sense (reverse complement of v-sense). The observed single-nucleotide reversion mutation (g2005a) is indicated in bold and another spontaneous mutation (c2006t) is indicated in red. The resulting amino acid replacements in the AC4 protein (L118R and S119P) are also indicated. **B.** CabLCV Rep protein domains and motifs, including iteron-related (IR) residues 7–11 (amino acids SFRLA) and the geminivirus Rep sequence (GRS; Nash et al. 2011). The coordinates of the oligomerization domain and RETINOBLASTOMA protein interaction surface are inferred based on alignment to the Rep protein sequence from tomato golden mosaic virus (Orozco and Hanley-Bowdoin 1998, Kong et al. 2000, Reyes 2012). Walker motifs were inferred based on Ruhel et al. (2021). Positions of amino acids L145 and E176 are shown with a purple line and a blue line.

In this work, we used the CabLCV-*A. thaliana* pathosystem for genome-wide analysis of spontaneous virus mutation, both in the presence and absence of a strong selective pressure. We built on previous work using infectious clones of CabLCV in which the Replication-associated gene (Rep) was mutagenized (Argüello-Astorga et al. 2007), extending analyses that demonstrated rapid single-nucleotide mutation in this mutagenized region. Such substitutions are thought to restore high-affinity interaction with the host RETINOBLASTOMA family protein required for reprogramming of DNA replication (Argüello-Astorga et al. 2004). Surprisingly, we identified additional convergent mutations in both virus segments (DNA-A and DNA-B) in a majority of inoculated plants. The allele frequency of these mutations varied with virus genotype (wild-type [WT] vs. mutagenized) and host genotype.

## Materials and Methods

Two *A. thaliana* accessions were used – the susceptible standard laboratory accession Columbia-0 (Col-0) and the highly susceptible natural inbred line Sei-0 (from Seis am Schlern, Italy; Kranz and Kirchheim 1987). Plants were grown at 20 °C with an 8 hours light/16 hours dark cycle in Percival Scientific plant growth chambers.

CabLCV infectious clones were previously described (Turnage et al. 2002). Plasmids pCPCbLCVA.003 and pCPCbLCVB.002 include partial tandem dimers of DNA-A and DNA-B. Resequencing these clones (see below) confirmed that the monomer unit segment sequences (2583 and 2512 nt, respectively) are identical to GenBank accessions U65529.2 and U65530.2 (Abouzid et al. 1992), with the exception of a single-nucleotide deletion adjacent to base 2368 (relative to the nick site) of DNA-B. Construction of mutagenized DNA-A plasmid pNSB1101 was previously described (Argüello-Astorga et al. 2007). Sequencing of that plasmid confirmed the presence of three introduced mutations and revealed a fourth, silent mutation in the Rep gene. Mutant alleles are illustrated in Figure 1A.

Biolistic inoculations at 21 days post-stratification were performed with a Microdrop Sprayer (Venganza, Inc). Inoculations used 2.5 μg of DNA per segment precipitated onto 1 μm gold particles (Cabrera-Ponce et al. 1997) and 30 psi for delivery. Plants were kept under a plastic lid for 24 hours after bombardment, then the lid was removed. Leaf tissue was collected 28 days post-inoculation (dpi) (Ascencio-Ibáñez et al. 2008). Tissue was flash frozen in liquid nitrogen and stored at −80 °C until further analysis.

DNA was extracted with cetyltrimethylammonium bromide (CTAB; Doyle and Doyle 1987) and subjected to PCR (for Sanger sequencing) or rolling-circle amplification with the GE TempliPhi kit (for Illumina sequencing) per the manufacturer's instructions. PCR primer sequences were FWD 5’-TCAGTCGTCAAACCCACATAC-3’ and REV 5’-CGAGGTGCCCACTCAAATAA-3’. PCR products were resolved on a 1% w/v agarose gel and purified using a QIAquick Gel Extraction kit (Qiagen, Hilden, Germany) according to the manufacturer’s instructions. Chromatograms were aligned to the reference sequence with Benchling (https://benchling.com) and are available at https://zenodo.org/record/5718844.

Nine libraries were prepared from phi29-amplified DNA with NEBNext index primers and the Ultra II kit (New England Biosciences, MA, USA). Two libraries were prepared from DNA-A plasmid inoculum (pCPCbLCVA.003 and pNSB1101). Multiplexed libraries were sequenced across two MiSeq flow cells with 150-bp paired-end reads. A DNA-B plasmid inoculum library was prepared at a later time with the Nextera XT kit and sequenced on a NovaSeq 6000 S4 lane (multiplexed with unique dual index primers). Illumina data were deposited in the NCBI Sequence Read Archive as PRJNA782339.

Illumina reads were trimmed with CutAdapt v1.16 (Martin 2011), aligned with BWA MEM v0.7.13 (Li 2013), and processed with samtools v1.8 (Li et al. 2009). Variants were called with VarScan v2.4.4 (Koboldt et al. 2012) and annotated with snpEff v4.3t (Cingolani et al. 2012). Variant calls and allele frequencies were analyzed with R 4.1.1 (R Core Team 2021; Wickham et al. 2019). Job scripts and R code are available at https://zenodo.org/record/5736525.

## Results

Rep reversion and global mutation detection

To analyze CabLCV mutation under different scenarios, we inoculated two *A. thaliana* genotypes with infectious clones of WT or mutagenized virus. The mutagenized form of the virus included four nucleotide substitutions (Figure 1A): three adjacent substitutions (a2005g, g2006c, a2007c) causing an L145A substitution in the Rep protein (Figure 1B) and a silent mutation at position 1911 in DNA-A (E176E in Rep). The three adjacent mutations also cause L118R and S119P amino acid substitutions in the overlapping AC4 coding sequence (Figure 1A), residues that do not have known functions. The experiment is shown schematically in Figure 2A and 2B.

**Figure 2:**
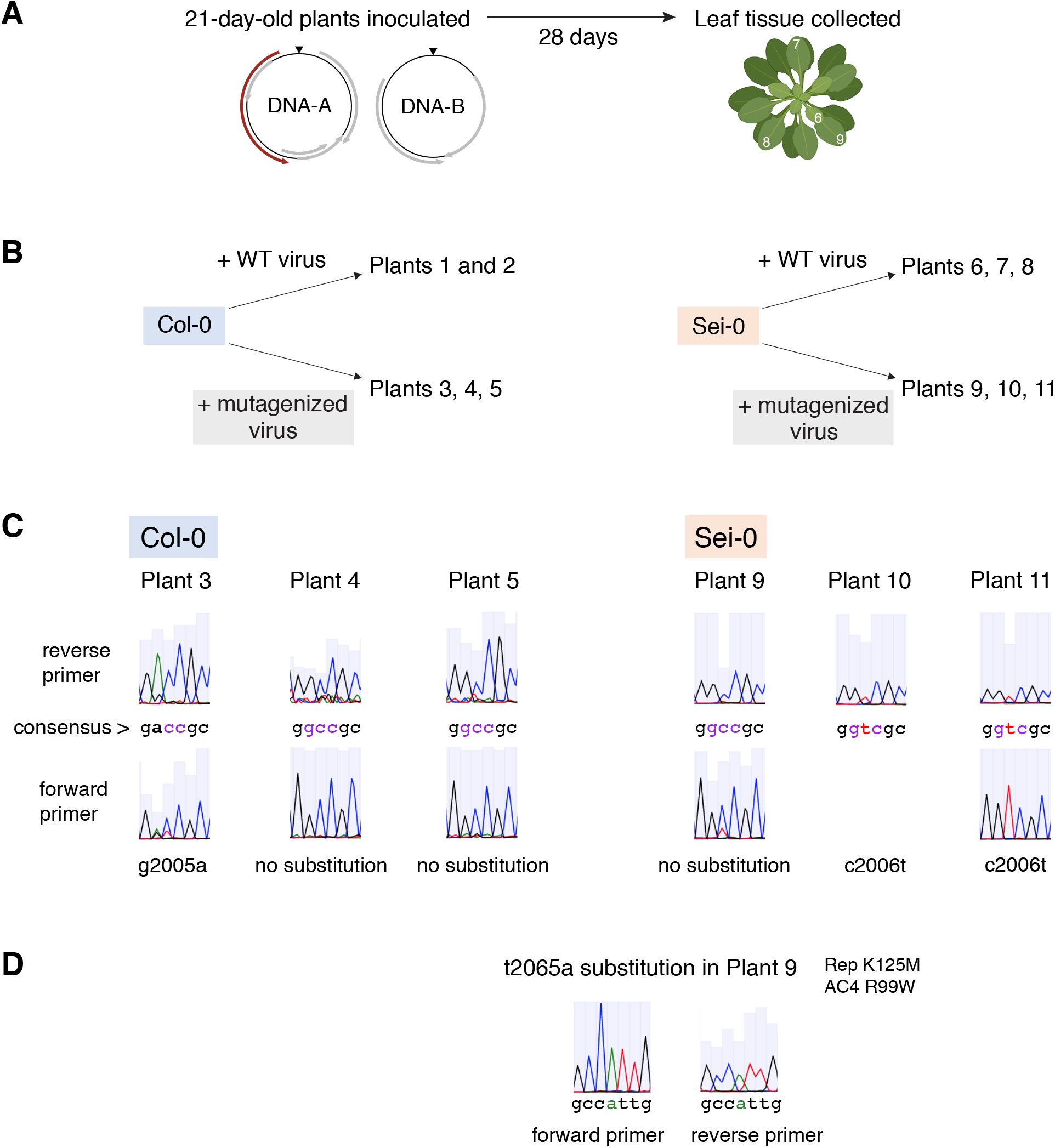
Rep reversion Sanger sequencing results. **A.** Experimental timeline. Leaves at approximate positions 6 to 9 from the center of the meristem (numbered in white) were pooled for DNA extraction. **B.** Structure of the experiment: two plant genotypes (Col-0 and Sei-0), each inoculated with two CabLCV genotypes. **C.** Sanger sequencing peaks showing the consensus sequence for six plants at 28 days after inoculation with mutagenized virus. (No forward primer chromatogram is shown for plant 10 because that reaction failed.) **D.** Sanger sequencing peaks for the one other consensus substitution (t2065a) detected in Rep gene amplicons. The *A. thaliana* rosette in panel A is from Biorender.com.

Rep PCR amplicons from eleven plants representing four experimental groups (Figure 2B) were characterized by Sanger sequencing. Consensus nucleotide substitutions were detected (by visual inspection of DNA sequence chromatograms) in three of the six plants infected with mutant virus (Figure 2C). Plant 3 (Col-0) showed direct reversion at DNA-A position 2005, resulting in a valine at Rep position 145, and plants 10 and 11 (Sei-0) had a mutation to thymine at position 2006 to give a threonine at Rep position 145 (Figure 2C). Both mutations were observed in a previous study (Argüello-Astorga et al. 2007) and are thought to partially restore interaction of Rep with the host RETINOBLASTOMA protein and increase virus replication. The consensus (mutagenized) sequence remained unchanged at these positions in the other half of the plants (#4, 5, and 9; Figure 2C). All six plants bombarded with mutagenized inoculum had consensus cytosine at position 1911, as expected based on the sequence of the inoculum plasmid (t1911c).

Only one other consensus substitution was evident in the Rep amplicon Sanger traces, i.e. a t2065a mutation in plant 9 (Figure 2D) causing a K125M substitution in the Rep protein. The possible fitness effects of this mutation are unclear given that it was only detected in a single plant. Based on its position in the protein (and the lack of a consensus L145 change in plant 9), it may affect interaction with RETINOBLASTOMA protein.

After rolling-circle-amplification, DNA from the same eleven plants was analyzed by deep sequencing to detect mutations genome-wide. Over 10,000 virus-mapping reads were obtained for both DNA-A and DNA-B for all plants, with two exceptions (low DNA-B read counts for Sei-0 plants 6 and 7). Most of the remaining reads mapped to the host reference genome (Supplementary Table S1). Read depth across DNA-A and DNA-B for each plant is shown in Supplemental Figure S1. We called variants (single-nucleotide replacements, small insertions, and small deletions) and, for simplicity, initially considered them as independent mutant alleles, not attempting to infer their local or global haplotype context.

No variants with estimated allele frequencies above 1% were identified for DNA-A in plants 6 or 7 or for DNA-B in plant 8 (three Sei-0 plants bombarded with WT virus) despite high read counts for those virus segments. Between 2 and 25 variants with an estimated frequency of at least 1% were detected for both segments in all other plants and were broadly distributed across the genome (Figure 3 and Supplementary Tables S2 and S3).

**Figure 3:**
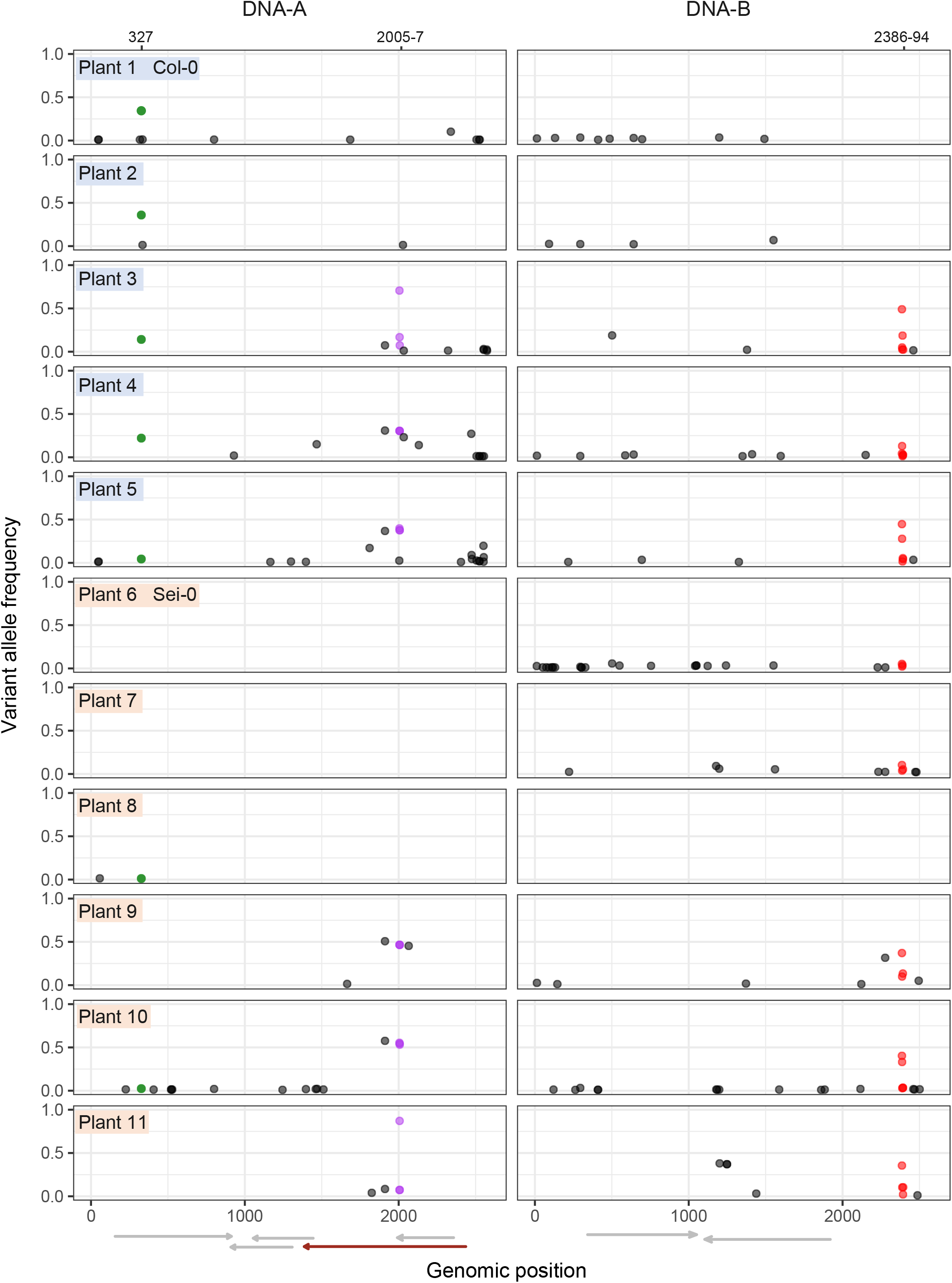
Variant allele frequencies for spontaneous mutations detected across DNA-A and DNA-B. Three recurrent mutation loci are highlighted: DNA-A g327a (coat protein S56N; green), Rep L145 reversion (purple) and a DNA-B mutational hotspot (red).

Majority variant calls from the Illumina data were generally concordant with the Sanger sequencing results shown in Figure 2. Subconsensus variants called in the mutagenized region also matched a minor dye peak for plant 3 (subconsensus c2006t, estimated at 16.8%). However, a puzzling pattern of highly correlated allele frequencies for variant calls corresponding to direct reversion of all four mutagenized positions was observed in the Illumina results without any corresponding evidence in the Sanger data. This pattern was particularly pronounced in plants 9 and 10 in which the nominal estimates of the WT nucleotides were all at ~50% (overlapping purple data points in

Figure 3; see Supplementary Table S2). Examination of individual reads showed the presence of all four substitutions in phase, indicating that all four were linked together in template molecules. Reversion at all four positions is hypothetically possible but would need to occur near-simultaneously for such linkage to occur. A more likely explanation is that this result represents a technical artifact caused by contamination with WT plasmid that affected all four positions equally. Based on the Sanger results, it is unlikely that contamination was present at the time of inoculation, especially given that WT virus would outcompete the mutagenized form. WT virus induces earlier symptoms than mutagenized virus (Argüello-Astorga et al. 2007), so the lack of such early symptoms for these plants strongly supports the idea that WT virus was not present. Circular plasmid DNA is an ideal substrate for phi29 polymerase (Dean et al. 2001) and would be enriched during rolling circle amplification. This interpretation indicates that the Illumina-based allele frequency estimates were conservatively biased, i.e. they represent lower bounds on the frequency of spontaneous mutations. Different plants inoculated with mutant virus appear to have been affected to differing extents, preventing full quantitative comparison. However, contamination would not impact the detection of medium-to-high abundance spontaneous mutation, allowing us to focus on the *presence* of these mutations for the remainder of this paper.

### Recurrent coat protein S56N substitution

The second most prominent feature in the DNA-A variant calling results was a mutation (g327a) detected in all five Col-0 plants and in two Sei-0 plants (#8 and 10, inoculated with WT and mutagenized virus, respectively; Figure 3). The resulting amino acid substitution (S56N) may disrupt phosphorylation of the viral coat protein (CP). The corresponding serine residue in the CP of African cassava mosaic virus (S62) is phosphorylated (Hipp et al. 2019). Repeated observation of this substitution strongly suggests that it is selectively beneficial under our experimental conditions, particularly in the two Col-0 plants inoculated with WT virus, which reached mutation frequencies of 34% and 36% (Figure 3).

### DNA-B iteron mutational hotspot

Illumina-based variant analysis revealed mutations across DNA-B in 10 of 11 plants, but no variants above 1% were detected in Sei-0 plant 8 (Figure 3). Most of the mutations detected across all plants (47 out of 77) map to the noncoding regions of DNA-B. Most mutations in the BV1 and BC1 coding regions were nonsynonymous (Supplementary Table S3).

The highest frequency DNA-B mutation observed across all plants was a g2386a mutation at 49.0% in plant 3 (a mutant-virus-inoculated Col-0 plant). This same mutation was observed in all six plants inoculated with the mutagenized virus and two of the three Sei-0 plants inoculated with WT virus (plants 6 and 7, at 5.3% and 10.4%, respectively). Mutations were also observed, albeit at lower frequency, at the adjacent nucleotide (g2387a) and the next few positions in the same plants (Figure 4). Thus, these mutations define a mutational hotspot overlapping with a candidate Rep-binding site (iteron). This site is in the region of highest sequence identity between DNA-A and DNA-B, i.e. the common region (Figure 4A). Mutations were also observed in DNA-A iterons, including a few that rose to high frequency (a2472c at 27% in plant 4, g2474t at 9% in plant 5), but not in such a pronounced cluster. The mutations in both DNA-A and DNA-B shifted the sequence away from the predicted optimum binding sequence of

**Figure 4:**
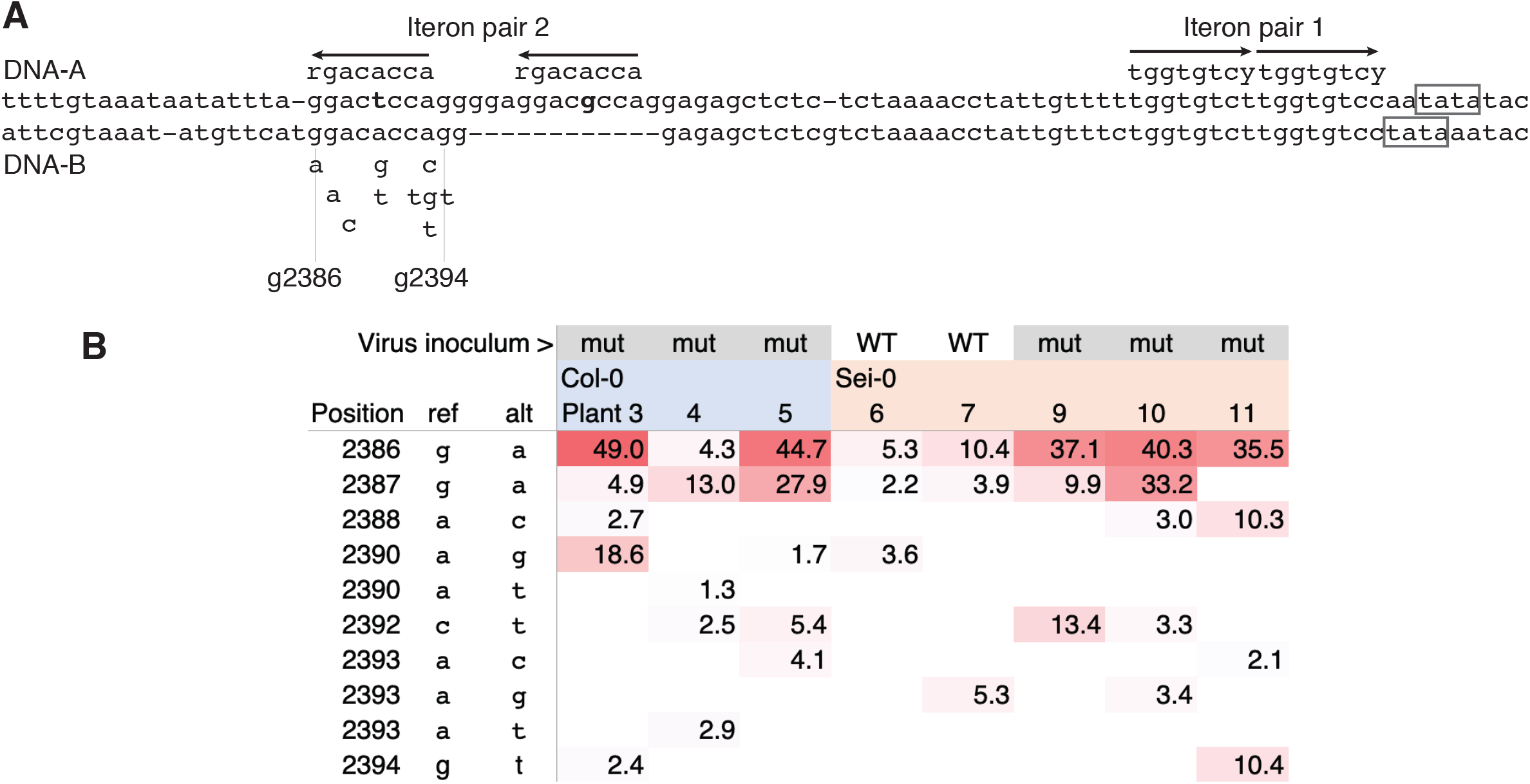
A mutational hotspot at a DNA-B iteron. **A.** Alignment of a section of the DNA-A/DNA-B common region showing two pairs of candidate Rep binding sites (iterons). The 8-nt presumed optimum binding sequence for CabLCV Rep is indicated (arrows) and two mismatches to this optimum in DNA-A are indicated in bold. The DNA-A region is positions 2439 to 2531, starting immediately adjacent to the AC1 (Rep) start codon, and the DNA-B region is positions 2368 to 2448. TATA boxes (reverse complement) for AC1 and BC1 are indicated. Several spontaneous mutations detected in DNA-B are indicated below that sequence (g2386a, g2387a, …). **B.** Heatmap table of estimated allele frequencies for mutations observed in the iteron. Mutations at these positions were not detected in plants 1, 2, or 8.

CabLCV Rep (tggtgtcy; Argüello-Astorga and Ruiz-Medrano 2001). Based on variant allele frequencies, selection on these positions was likely stronger in plants inoculated with mutagenized virus than in those inoculated with WT virus. The functional basis for the adaptive benefit of these mutations is not immediately clear, as discussed below.

## Discussion

We have documented rapid and reproducible spontaneous mutations in three regions of the CabLCV genome, one engineered (Rep helix 4) and two unanticipated (CP S56 and a DNA-B iteron). High-throughput sequencing revealed subconsensus variants across the genome, but the degree to which such spontaneous mutations are selectively advantageous is unknown. Future studies could benefit from increased biological and technical replication, particularly because detectable contamination with infectious clones limited our ability to make quantitative comparisons of mutant allele frequencies. We saw no obvious patterns of variant co-occurrence across plant samples indicating sample-to-sample cross-contamination with actual virus DNA (Supplementary Tables S2 to S5) but including additional controls (spike-in sequence identifiers or mock-inoculated plant DNA libraries) could rule out or mitigate this possibility. Higher sensitivity for rare variants could potentially be achieved with alternative DNA preparation strategies (Aimone et al. 2021b; Pinto et al. 2021).

### Implications for Rep-DNA and Rep-protein interactions

Although mutagenized CabLCV used here and in the previous study by Argüello-Astorga et al. (2007) is an artificial system, consistent results were obtained with a geminivirus from a different genus, maize streak virus (Shepherd et al. 2005). Similar to the K125M substitution observed here (Figure 2D), Argüello-Astorga et al. detected a substitution (I167L) on the other side of the RETINOBLASTOMA protein interaction region. It is unclear whether these second-site substitutions directly compensate for disruption of the protein-protein interaction, but these results suggest that epistatic interactions among Rep codons should be explored further.

The strong selective sweep for restored Rep function did not prevent selection on other loci but may have weakened selection for the CP S56N substitution in Col-0 plants. By contrast, the use of mutagenized inoculum appears to have enhanced selection for mutant alleles at the DNA-B iteron, particularly for the g2386a substitution (Figure 4). This iteron may be associated with one or more Rep-mediated functions, including in DNA-B replication, transcriptional repression, or other as-yet-unknown molecular interactions. Rep auto-represses its own transcription via a DNA-A iteron (Haley et al. 1992; Hanley-Bowdoin et al. 1999), though no similar function has been described for a corresponding DNA-B iteron. Alternatively, the normal function of this region may be unrelated to Rep binding and/or the region may have no molecular function at all. CabLCV has an atypical Rep structure and the CabLCV DNA-A/DNA-B common regions have a greater number of differences than is typical for begomoviruses (Hill et al. 1998), making it difficult to extrapolate from better studied viruses such as tomato golden mosaic virus (Orozco et al. 1998). Experimental assessment of the function of this iteron could clarify why these mutations increase fitness. It may be that the substitutions disrupt Rep binding to DNA-B, reducing competitive sequestration of Rep molecules and thus enabling better DNA-A replication. If Rep binding to this DNA-B site is less important, increased DNA-A replication could indirectly provide a fitness benefit to DNA-B. We would expect the Rep L145A substitution, which impairs replication of both molecules, to enhance this benefit, consistent with our observations.

The DNA-B long intergenic region was also a mutational hotspot during experimental evolution in cassava (Aimone et al. 2021c), suggesting this pattern will occur in other experimental and natural systems.

### CP and host-genotype-specific differences in selection pressure

The CP gene is dispensable for infection in many laboratory situations (Stanley and Townsend 1986; Pooma et al. 1996; Turnage et al. 2002) so strong selection on a CP substitution was not necessarily expected. This result highlights the use of experimental evolution as an efficient tool for ‘forward’ genetic screening (Cooper 2018). Similar to what we have found here, an ssDNA circovirus also displayed high mutation frequencies within the coat protein gene (Correa-Fiz et al. 2020). Similar results were also observed for tomato leaf curl begomoviruses (Sánchez-Campos et al. 2018). Sequencing virus populations from experimental inoculation studies may become a routine step in characterizing new infectious clones as sequencing costs decline. Introducing mutations that rapidly rise in frequency back into their ‘parent’ infectious clones may reduce experimental variability due to possible phenotypic effects from these mutations.

The S56N substitution detected in 7 of the 11 plants in this experiment likely prevents phosphorylation, providing a possible explanation for its selective benefit. Hipp et al. (2019) detected by mass spectrometry partial phosphorylation of three N-terminal residues (one homologous to the CabLCV residue) in the CP of African cassava mosaic virus. Hipp et al. suggested that this phosphorylation may promote ubiquitin-dependent proteasomal degradation of CP, similar to degradation of tomato yellow leaf curl virus CP (Gorovits et al. 2014, 2016). The S56N substitution may stabilize CP, enhancing its (largely unknown) functions in cell-to-cell and long-distance movement. The N terminus of CP also functions in nuclear localization and in ssDNA binding during virion assembly (Unseld et al. 2001, 2004), so other effects remain possible. This amino acid position is conserved across many geminiviruses, with the serine sometimes replaced with threonine (Hipp et al. 2019), another residue that can be phosphorylated. Therefore, it is possible that the within-plant selective advantage we have observed is counterbalanced by long-term costs, particularly if the substitution interferes with packaging and/or whitefly transmission.

The basis for the stronger effect for S56N in Col-0 relative to Sei-0 is not clear, but should be tractable for future study. The geminivirus beet curly top virus also causes strong symptoms on Sei-0, likely because it accumulates to high levels (Park et al. 2004; Lee et al. 1994), but differences in DNA-A and DNA-B levels in Col-0 vs. Sei-0 were variable in our experiment (Supplemental Table S1). Sei-0 can be infected with African cassava mosaic virus, whereas Col-0 cannot (Aimone et al. 2021a). The genetic basis for the hypersusceptibility of Sei-0 is currently unknown but could be genetically mapped.

*A. thaliana* can be infected with a number of geminiviruses (Ouibrahim and Caranta 2013), but it is unclear which, if any, of these viruses are well-adapted to it. Serially passaging CabLCV in *A. thaliana* would presumably lead to further substitutions, some of which may involve trade-offs including reduced ability to adapt to changing conditions, as observed for turnip mosaic virus (Butković et al. 2020; González et al. 2019). CabLCV is thought to be an ecologically realistic begomovirus for challenging *A. thaliana* based on its broad host range within the Brassicaceae and Fabaceae (Strandberg et al. 1991; Fiallo-Olivé et al. 2018), but this assumption should be reexamined. Measuring the prevalence of geminiviruses in natural *A. thaliana* populations, as has been done for several RNA viruses (Pagán et al. 2010), would be informative and further enhance the value of this experimental system.

## Acknowledgements

We thank the Office of Advanced Research Computing (OARC) at Rutgers, The State University of New Jersey, for maintaining and providing access to the Amarel cluster. OWW, EW, SR, AM, and KRHM worked under the NCSU Biochemistry Undergraduate Research and Training Program (BURT-P). EW, SR, AM, and KRHM were partially supported by the T & E Biochemistry Fund. We thank David O. Deppong and Mary M. Dallas for laboratory assistance. We thank Linda Hanley-Bowdoin and Alvin Crespo-Bellido for helpful discussions and comments on the manuscript.

## Author contributions

JTAI designed the project. OWW, AM, EW, SR, KRHM, AED and JTAI performed experiments. JSH and IC processed Illumina data. JSH and JTAI drafted the manuscript. All authors analyzed data and revised the manuscript.

---

**Figure S1:**
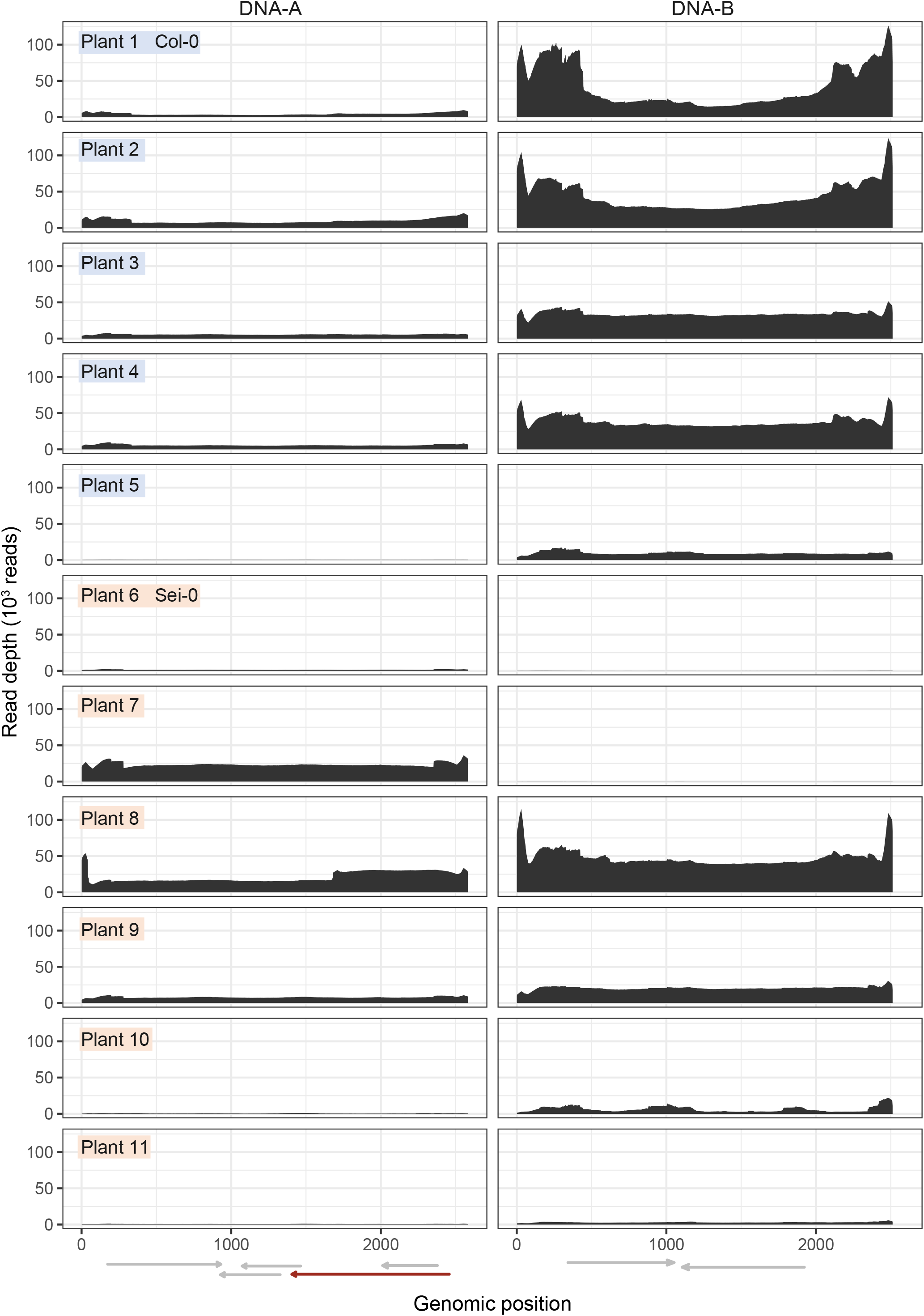
Illumina read depth across virus segments DNA-A and DNA-B.

## Notes

https://www.ncbi.nlm.nih.gov/sra/PRJNA782339

https://zenodo.org/record/5718844

https://zenodo.org/record/5736525

### Competing Interest Statement

The authors have declared no competing interest.

### Summary of Updates

A few sentences added or updated in the Introduction, Results, and Discussion sections to improve clarity. Add supplemental graphs of read depth.

